# Genome-wide association and genomic prediction of resistance to *Flavobacterium columnare* in a farmed rainbow trout population

**DOI:** 10.1101/2022.02.28.482244

**Authors:** Clémence Fraslin, Heikki Koskinen, Antti Nousianen, Ross D. Houston, Antti Kause

## Abstract

Columnaris disease is an emerging disease affecting farmed rainbow trout (*Oncorhynchus mykiss*) globally. In aquaculture breeding, genomic selection has been increasingly used to improve traits that are difficult to measure on candidate fish (such as disease resistance traits). Following a natural outbreak of columnaris disease, 3,054 exposed fish and their 81 parents (33 dams and 48 sires) were genotyped with the 57K SNP Axiom™ trout genotyping array. Genetic parameters of host resistance (measured as a binary survival trait) were estimated, a genome wide association study was performed, and the accuracy of pedigree-based and genomic prediction was estimated. After quality controls, 2,874 challenged fish (1,403 dead fish and 1,471 alive fish) and 78 parents genotyped for 27,907 SNPs remained. Pedigree based heritability was estimated to be 0.18 and 0.35 on the observed and underlying scale, respectively. Genomic heritability was estimated to be 0.21 and 0.43 on the observed and underlying scale, respectively. A quantitative trait loci (QTL) was detected on chromosome Omy3, significant at the genome-wide level, along with several suggestive QTLs on two other chromosomes. The additive effect on mortality proportion of the peak SNP from Omy3 was estimated to be 0.11 (0.018; se). Pedigree-based prediction accuracy was 0.59, and the use of genomic evaluation increased the prediction accuracy by at least 13.6%. Using the second iteration of a weighted genomic-based evaluation increased the prediction accuracy by 18.6% compared to the pedigree-based model. These results suggest that resistance is a suitable target trait for genetic improvement by selective breeding, and genomic selection may be a useful approach to speed up this process.

## 1. Introduction

Rainbow trout (*Oncorhynchus mykiss*) is an important aquaculture species globally, and is produced both in freshwater and sea water. Infectious diseases are a major threat to aquaculture production worldwide, with major impacts on fish welfare, the environment, and the sustainability of aquaculture (Houston, 2017; Yáñez et al., 2014). Water temperatures are rising due to global warming, and fish farms could be subject to more frequent and longer periods of warm water, conditions often more conducive to fish pathogens (Karvonen et al., 2010). One of the pathogens that take advantage of warmer temperature is *Flavobacterium columnare*, a gram-negative bacterium, responsible for columnaris disease (CD) that affects various fish species of importance for aquaculture worldwide, including rainbow trout (Declercq et al., 2013). Outbreaks of CD occur in rainbow trout farms mainly during the summer, when the water temperature rises above 18°C, with up to 100% mortality in the absence of antibiotic treatment (Pulkkinen et al., 2010; Suomalainen et al., 2005a). *F. columnare* causes both acute and chronic infections with necrosis of tissues resulting in skin lesions, fin erosions, mouth rot and gill necrosis often leading to the death of the fish (Declercq et al., 2013). A modified live *F. columnare* vaccine for channel catfish (*Ictalurus punctatus*) and largemouth bass (*Micropterus salmoides*) has been developed and tested (Shoemaker et al., 2011) and is now available commercially AQUAVAC-COL™. However, for rainbow trout there is no licensed commercial vaccine available against *F. columnare.* The treatments used in rainbow trout consist of adding salted water to increase the salinity (Declercq et al., 2013; Suomalainen et al., 2005b) or using antibiotics, either in a bath treatment or in the feed (Bullock, 1986). The use of antibiotics and antimicrobial agents to treat CD is not a sustainable solution as it can contribute to antimicrobial resistance, a major concern for human and animal health (Serrano, 2005). Thus, other means to control the disease are needed.

Increasing innate genetic resistance of rainbow trout through selective breeding could be a sustainable solution to this major problem. The Finnish national breeding programme for rainbow trout was established in the late 1980’s, targeting production traits such as growth, age of maturity, external appearance, fish welfare, visceral percentage, and survival recorded on fish reared in brackish sea and fresh water (Kause et al., 2003; Kuukka-Anttila et al., 2010; Vehviläinen et al., 2012). To date, estimation of breeding values has been based on pedigree information that is obtained by rearing families initially separated in a large number of family tanks followed by individual tagging and pooling of all the fish. The development and availability of new genomics tools such as genotyping-by-sequencing or single nucleotide polymorphism (SNP) arrays (Palti et al., 2015; Robledo et al., 2017) have facilitated studies of the genetic architecture of valuable production traits. The same technologies have also supported the testing and implementation of genomic selection (GS) in aquaculture breeding programmes for several major species for aquaculture (Houston et al., 2020; You et al., 2020). Genomic selection relies on the use of thousands of genetic markers (such as SNPs) spread over the entire genome to estimate the breeding values of selection candidates. A reference population with both phenotypes and genotype data is used to train a prediction model that is then used to estimate breeding values of the candidates that are typically genotyped but not phenotyped (Meuwissen et al., 2001). Although genomic selection has been routinely implemented commercially in the major aquaculture species such as Atlantic salmon (Norris, 2017), it is still in its early days for other species. Recent studies show that in rainbow trout, resistance against *F. columnare* is indeed heritable and that genomic selection could be a potential way to improve the resistance (Evenhuis et al., 2015; Silva et al., 2019a, 2019b).

The objective of this study was to investigate the genetic architecture of resistance to *F. columnare* and quantify the potential of genomic selection to improve resistance in a rainbow trout breeding programme. Specifically, the heritability of resistance to *F. columnare* was estimated in a rainbow trout population from Finland, then a Genome Wide Association Study (GWAS) was performed to investigate the genetic architecture of resistance, and finally, the accuracy of breeding value predictions was compared between genomic evaluation and traditional pedigree-based evaluation approaches. The results contribute to the cumulating evidence of the benefits and suitable ways of implementing genomic selection in aquaculture breeding.

## 2. Material and Methods

### 2.1. Ethical statement

The establishment of progeny families at Luke's research facilities followed the protocols approved by the Luke's Animal Care Committee, Helsinki, Finland. Hanka-Taimen Oy, a fish farming company, has authorisation for fish rearing and experiments, and both parties comply with the EU Directive 2010/63/EU for animal experiments.

### 2.2. Fish rearing and phenotyping

On the 15^th^ of May 2019, 105 rainbow trout families (from 33 dams and 48 sires) were produced from the Finnish national breeding programme maintained by Luke at Enonkoski research station in east Finland. After stripping and fertilisation, a sample of eggs from all families were pooled and sent to the multiplier farm of Hanka-Taimen Oy (Hankasalmi, Finland). The eggs were incubated and fingerlings reared in bulk. In June 2019, around 30,000 fry were distributed into three fingerling tanks, about 100 fish per family per tank and the fish were reared following commercial practices.

The multiplier farm uses water from a nearby stream, and natural *F. columnare* outbreaks occur frequently. From the day the fish were in the three tanks (day 0 of the study), signs of any disease and mortality were monitored twice a day. On days 11 and 14 to 16, fish in all three tanks started to show signs of CD. From day 20 to 24 a piece of tail was taken from around 510 fish per tank, randomly chosen amongst the dead fish with clear CD signs, for later genotyping. These 1,538 fish were considered as susceptible fish in the analysis. Some of the fish with clear CD sings were sent for veterinary examination, and after CD infection was confirmed by a vet from sampled dead fish, the three tanks were treated for *F. columnare* with approved treatments of salt, chloramine, and medical feeds. CD signs and increased mortality were observed also on days 56 and 71 and fish were treated again according to an approved treatment. On the last day of the study (day 99, October 2019), a piece of tail was collected, for later genotyping, on around 506 alive randomly sampled fish per tank. These 1,519 fish were considered as resistant fish. During the three months of the trial, water temperature was recorded daily.

### 2.3. Genotyping, quality controls, parentage and sex assignment

Samples from 3,057 fish from the trial and 567 fish from the parental generation including the 81 parents were all sent to IdentiGEN Ltd. (Dublin, Ireland) for DNA extraction and genotyping using the 57K SNP Axiom™ Trout Genotyping Array (Palti et al., 2015). Prior to the calling of genotypes, quality controls on the 57,501 SNPs from the SNP array were performed following D’Ambrosio et al., (2019). The SNP probes were aligned, using a BLASTn® procedure to the Omyk_1.0 genome assembly available at the time of the analysis (Gao et al., 2018; Pearse et al., 2019). Only SNPs with a unique position on the genome (50,820 SNPs) were retained. Genotypes of all 3,621 sampled fish were called together in a single run using Axiom Analysis Suite software (v. 4.0.3.3) with standard SNP quality control (QC) thresholds and customized sample QC thresholds (DQC ≥ 0.82, QC call rate ≥ 90, percent of passing samples ≥ 80, average call rate for passing samples ≥ 95). SNPs that were classified as off target variant or other, monomorphic SNPs and SNPs for which no homozygous individual was observed for the minor allele were discarded and hence 36,020 polymorphic SNPs were kept for further analysis. Using plink software (v.1.9, Chang et al., 2015), further quality controls were performed on both SNPs and individuals. Duplicated individuals were detected using the --genome option from plink, two individuals with an identity-by-descent value over 0.90 were considered as duplicated and both individuals were removed from the analysis. Only the individuals with a call rate over 0.90 and the SNPs with a minor allele frequency (MAF) higher than 0.05, call rate higher than 0.95 and that passed the Hardy-Weinberg equilibrium test (*p*-value < 10^−6^) were retained. The final dataset comprised 3,435 fish (2,874 challenged fish, and 561 fish from the parental generation including 78 parents) genotyped for 27,907 high-quality SNPs.

Since three parents (two dams and one sire) were missing from the final dataset, parentage assignment was performed in two steps. First using a subset of 200 SNPs with a 100% call rate in both generations, the pedigree was reconstructed for fish with no missing parents using the R package APIS (Griot et al., 2020) with a mismatch assignment set to 1%. In the second step, the genomic relationship values derived from the genomic relationship matrix (GRM) built with GCTA software (Eq.1, Yang et al., 2011) were used to recover the family where one parent was missing from the genotyped dataset. The full pedigree was recovered for 97.6% of the fish while the remaining 68 fish had one parent unassigned / missing.

### 2.4. Genomic relationship matrix

The GCTA software was used to compute the GRM, and the genetic relationship between individuals j and k (*g*_jk_) was estimated following:

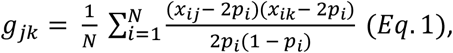

in which *N* is the total number of SNPs (27,907), *x*_*ij*_ and *x*_*ik*_ are the number of copies of the reference allele for the i^th^ SNP for the j^th^ and k^th^ fish, respectively, and *p*_*i*_ is the frequency of the reference allele estimated from the markers.

### 2.5. Estimation of genetic parameters

Variance components and heritability were estimated based either on pedigree-based relationships or GRM using ASReml (v.4.1, Gilmour et al., 2015) with two different approaches, a linear mixed model (Eq.2) and a logistic regression model (Eq. 3) to assess the trait on the observed and underlying scales, respectively:

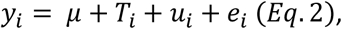

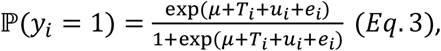

in which, for the i^th^ fish, *y*_*i*_ is the phenotype recorded as binary survival (0 for alive and 1 for dead fish), *μ* is the population mean, *T*_*i*_ is the fixed effect of tank (3 levels), *u*_*i*_ is the random additive genetic value of individual i, following a normal distribution 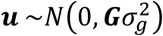 or 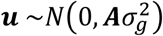 with 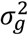 the estimated genetic variance, where **G** is the GRM built with GCTA (Eq.1) and **A** is the pedigree-based relationship matrix. Finally, *e*_*i*_ is the residual effect following a normal and independent distribution 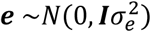 with 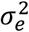 being the residual variance.

### 2.6. Genome wide association study

To identify SNPs associated with resistance to *F. columnare*, a GWAS was performed using a mixed linear model association (Eq.4) with the leave-one-chromosome-out (loco) option implemented in GCTA:

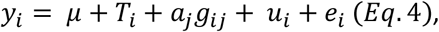

in which *y*_*i*_ is the observed phenotype of the i^th^ individual (0 for alive and 1 for dead), *μ* the overall mean in the population, *T*_*i*_ the fixed effect of the tank (3 levels), *a*_*j*_ is the additive genetic effect of the reference allele for the j^th^ SNP with its genotype for individual i (*g*_*ij*_) coded as 0, 1 or 2. And *e*_*i*_ is the residual effect following a normal and independent distribution 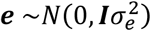 with 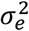 the residual variance. Finally, *u*_*i*_ the random vector of polygenic effects followed a normal distribution 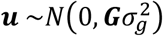 with 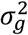 the estimated genetic variance and **G** a partial GRM constructed with 28 chromosomes after removing the chromosome containing the j^th^ SNP since the analysis was performed using the leave-one-chromosome-out (mlma-loco) approach.

### 2.7. QTL characterisation

For the GWAS, a Bonferroni correction with *α*=5% was used to determine the genome-wide significance threshold [−log_10_(*α*/*n*)] and the chromosome-wide suggestive threshold [−log_10_(*α*/[*n*/29])], with *n* the number of SNPs in the analysis. Only the SNPs with a −log10(*p*-value) over the chromosome wide threshold were considered to detect QTL associated with the resistance. For each QTL, the additive effect (*a*) of the top SNP was used to estimate the proportion of genetic variance explained by this peak SNP using:

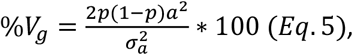

with 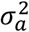 the total genetic variance estimated using the linear model (Eq.2) with ASReml and *p* the minor allele frequency of the target SNP.

Candidate genes located within a 2 Mb window around the peak SNP (1 Mb each side) for each QTL were investigated using the NCBI *Oncorhynchus mykiss* Annotation Release 100 (GCF_002163495.1).

### 2.8. Genomic prediction of breeding values

The efficiency of genomic prediction of breeding values was assessed using 20 replicates of Monte-Carlo “leave-one-group-out” cross-validation tests. For each replicate, fish were randomly assigned to five groups, four-fifths of the fish (*n*=2,300) with known phenotypes and genotypes were used as the training set and one fifth of the fish (*n*=575) with known genotypes and masked phenotypes were used as the validation set. Mixed linear BLUP animal model (Eq.2) and logistic regression model (Eq.3) were used to estimate pedigree-based (EBV) and genomic breeding values (GEBV) of fish in the validation set using phenotypic values of fish in the training set and the relationship matrix, based on pedigree or genomic information, using two different software, BLUPF90 (Misztal et al., 2002) and ASReml.

For the Genomic BLUP (GBLUP), two approaches were implemented using the BLUPF90 software, a standard GBLUP and a weighted GBLUP (wGBLUP) approach. The wGBLUP is an iterative approach in which, at a given k^th^ iteration, a weight is determined for each SNP based on the SNP variance (derived from the SNP additive effect) estimated at the (k-1) iteration (Wang et al., 2012). For the first iteration, weights are fixed to 1 which is equivalent to a standard GBLUP and we performed three iterations (labelled w2GBLUP and w3GBLUP).

The accuracy (*r*) of genomic prediction was computed as the Pearson correlation coefficient between the (G)EBV and the true phenotype (*y*) of the fish in the validation set divided by the square root of the genomic based heritability obtained on the observed scale (*h*^*2*^_*obs*_), following Legarra et al., (2008) using the formula:

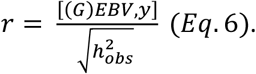

Since resistance was measured as binary survival, receiver operator characteristic (ROC) curves were also used to assess the accuracy of classifying animals as susceptible or resistant using the different models. The area under the ROC curve (AUC) metric was used to assess the performance of the classifier: an AUC value of 1 represents a perfect classifier and an AUC value of 0.5 representing a random classifier (Hanley and McNeil, 1982; Wray et al., 2010).

## 3. Results

### 3.1. Sample and data collection from farm outbreak

A small number of mortalities occurred in tanks 1 and 2 in the first few days after the transfer, with a peak at day 3 with 44 and 58 mortalities recorded in tank 1 and 2, respectively. Higher mortality was recorded after day 1 in tank 3, with a peak on day 4 (119 dead fish) after which mortality decreased to a base level of 30 mortality/day on average for a week. No specific causes were observed for these mortalities.

Mortalities accompanied with clear signs of *F. columnaris* started on day 16 for fish in tanks 1 and 2 respectively, and on day 18 for fish in tank 3 (Figure 1). On day zero at the end of June, the average daily water temperature was above 17°C and started to rise above 18°C from day 2 onwards with a peak temperature at 25°C on days 33 and 34. At the end of the recording period, the final mortality was 39.4% (Figure 1).

**Fig. 1.**
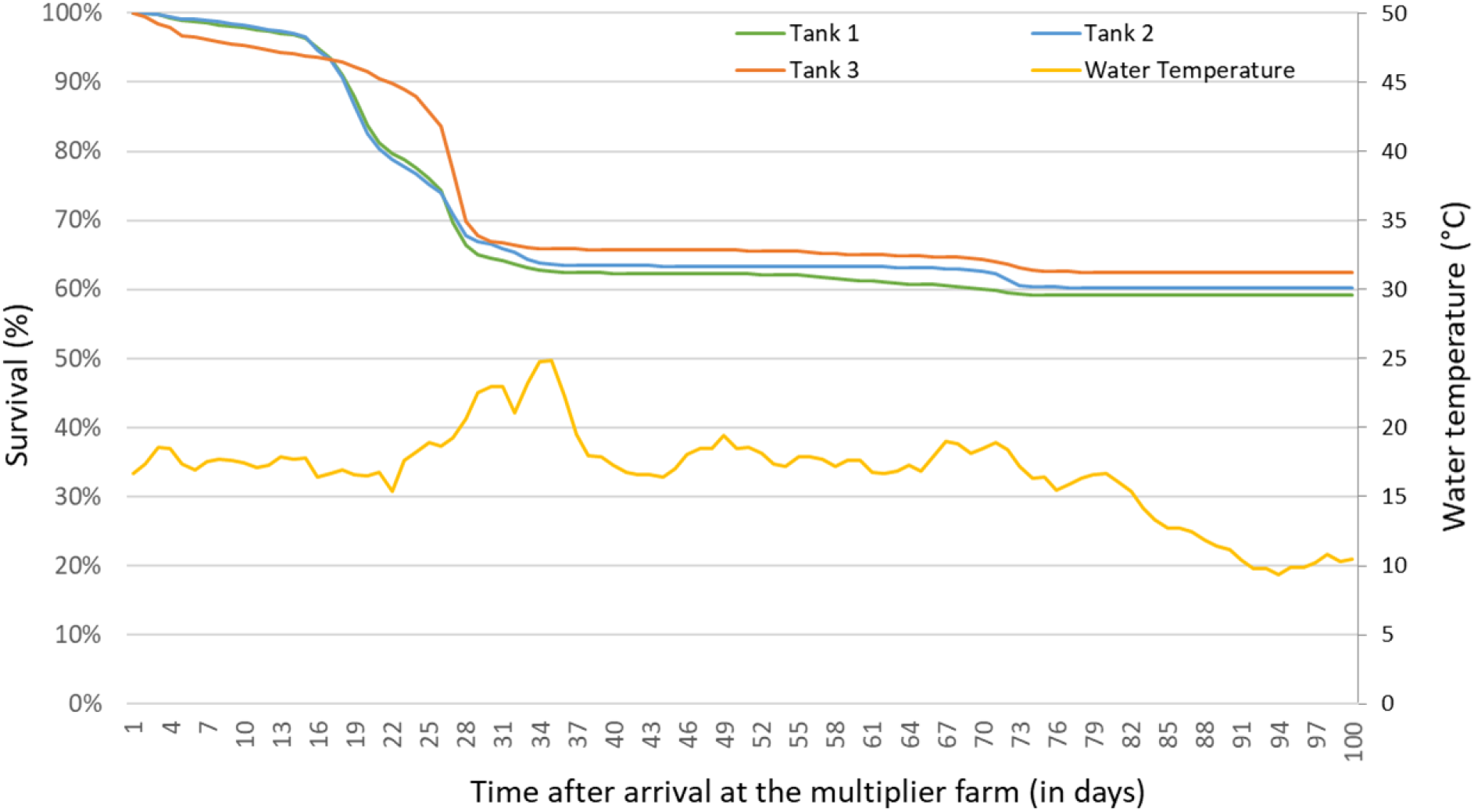
Survival curves and water temperature for the three tanks for the duration of the study at the multiplier farm.

### 3.2. Genetic parameter estimates

The estimates of genetic parameters using the linear and the logistic regression models are summarised in Table 1. Estimates of the pedigree-based heritability of binary survival were 0.18 (± 0.038; se) and 0.35 (± 0.046; se) on the observed and underlying scale, respectively. Genomic heritability of binary survival to *F. columnaris* were slightly higher than the pedigree-based estimates, at 0.21 (± 0.030; se) and 0.43 (± 0.042; se) on the observed and underlying scale, respectively.

**Table 1.** Estimates of variance components for resistance to *F. columnare*.

| Model | Relationship matrix | $\sigma_a^2 (\pm \text{se})$ | $\sigma_p^2 (\pm \text{se})$ | $\sigma_e^2 (\pm \text{se})$ | $h^2 (\pm \text{se})$ |
| --- | --- | --- | --- | --- | --- |
| Linear | A | 0.044 ± 0.010 | 0.25 ± 0.008 | 0.21 ± 0.008 | 0.18 ± 0.038<br>(0.34) |
| Linear | G | 0.054 ± 0.009 | 0.25 ± 0.008 | 0.20 ± 0.007 | 0.21 ± 0.030<br>(0.40) |
| Logit | A | 0.54 ± 0.108 | 1.54 ± 0.108 | 1 | 0.35 ± 0.046 |
| Logit | G | 0.76 ± 0.132 | 1.76 ± 0.132 | 1 | 0.43 ± 0.042 |

A: the pedigree-based relationship matrix
G: the genomic relationship matrix
Logit: logistic regression.
$\sigma_a^2$ : genetic variance.
$\sigma_p^2$ : phenotypic variance.
$\sigma_e^2$ : residual variance.
$h^2$ : For the linear models, heritability is estimated on the observe scale and transformed on the underlying scale (value in brackets) using the formula from Dempster and Lerner (1950). For the logistic regression models, heritability is estimated on the underlying scale with the residual variance fixed to 1 by convention.

### 3.3. Genome wide association study

A highly significant QTL affecting the binary trait of resistance was detected on chromosome 3 (Figure 2). There were a total of 28 SNPs that were significantly associated with resistance to *F. columnare*, with a −log_10_(*p*-value) that was over the 5% chromosome-wide Bonferroni threshold (−log_10_(*p*-value) = 4.28). Those SNPs were located within three chromosomes with 23 SNPs on Omy3, one SNP on Omy12 and four SNPs on Omy15 (Figure 2).

**Fig. 2.**
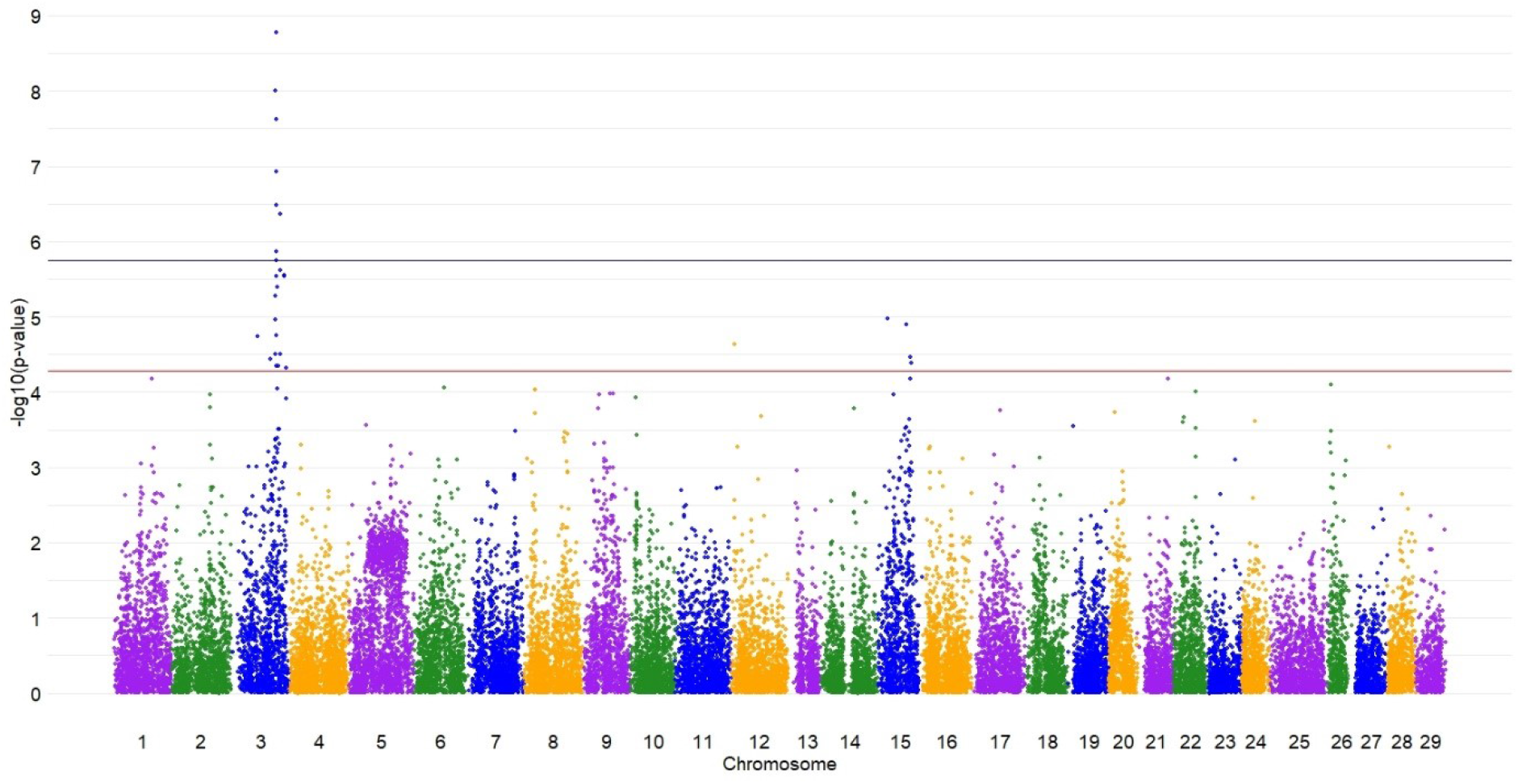
Manhattan plot of QTL associated with resistance to *Flavobacterium columnare* detected using a GWAS under a MLMA-LOCO model. The dark blue line is the 5% significance threshold at the genome-wide level, the red line is the 5% suggestive threshold at the chromosome-wide level.

On chromosome 3, 23 SNPs (one SNP at 38.165 Mb and the rest spanning from 55.715 Mb to 79.557 Mb) had a −log_10_(p-value) at least over the 5% chromosome-wide threshold level (Supplementary Table S1), including 8 SNPs with a −log_10_(p-value) that surpassed the 5% genome-wide threshold with the peak SNP located at 64.39 Mb (Table 2). The QTL explained an estimated 11.2% of the additive genetic variation and the additive effect of the peak SNP from Omy3 on mortality proportion (the binary resistance trait) was estimated to be 0.11 (0.0018; se), implying approximately 22% difference in survival between alternate homozygous fish at the QTL.

**Table 2.** Detection of QTLs associated with resistance to *Flavobacterium columnare* in rainbow trout.

| Chr | Peak SNP<br>name | Peak SNP<br>position (bp) | Peak SNP additive<br>effect (± se) | Peak SNP<br>*p*-value | Peak SNP<br>-log<sub>10</sub>(*p*-value) | %Vg explained<br>by peak SNP |
| --- | --- | --- | --- | --- | --- | --- |
| 3 | AX-89933339 | 64,390,419 | -0.11 ± 0.0018 | 1.69E-09 | 8.77 | 11.20% |
| 12 | AX-89951575 | 5,316,265 | -0.073 ± 0.0172 | 2.30E-05 | 4.64 | 4.40% |
| 15 | AX-89956130 | 41,172,606 | 0.093 ± 0.0213 | 1.25E-05 | 4.90 | 5.70% |

Chr: chromosome.
%Vg = $2p(1-p)a^2/0.054$ (Eq. 5), the proportion of total genetic variance explained by the peak SNP, with *p* the SNP minor allele frequency and *a* the SNP additive effect.

On chromosome 12, the only SNP that excessed the 5% suggestive threshold was located at 5.316 Mb and explained 4.40% of the total genetic variance of resistance to *F. columnare* (Table 2 and Supplementary Table S1).s On chromosome 15, the first suggestive SNP was located at 13.36 Mb and the remaining three significant SNPs spanned from 41.173 to 47.018 Mb with the peak SNP explaining 5.70% of the total genetic variance (Table 2).

The observed *p*-values were inflated with a *λ* value of 1.362 as expected for a population with large full and half-sib families (a *λ* of ^~^1.1 indicates a relatively good concordance between observed and predicted *p*-values, Yang et al., 2011b).

### 3.4. Genomic prediction of breeding values

For both linear and logistic models, genomic-based predictions of breeding values resulted in a higher accuracy than pedigree-based prediction, with an increase of 15.3% between the w3GBLUP and the pedigree-based BLUP (PBLUP) models and of 18.6% between the w2GBLUP and the PBLUP models (Table 3). Correlations and accuracies obtained under the linear or logistic regression models were very similar. The increase in accuracy between the pedigree-based and the genomic-based logistic regression (+13.6%) was slightly lower than using the genomic-based linear model (+15.3%).

**Table 3.**
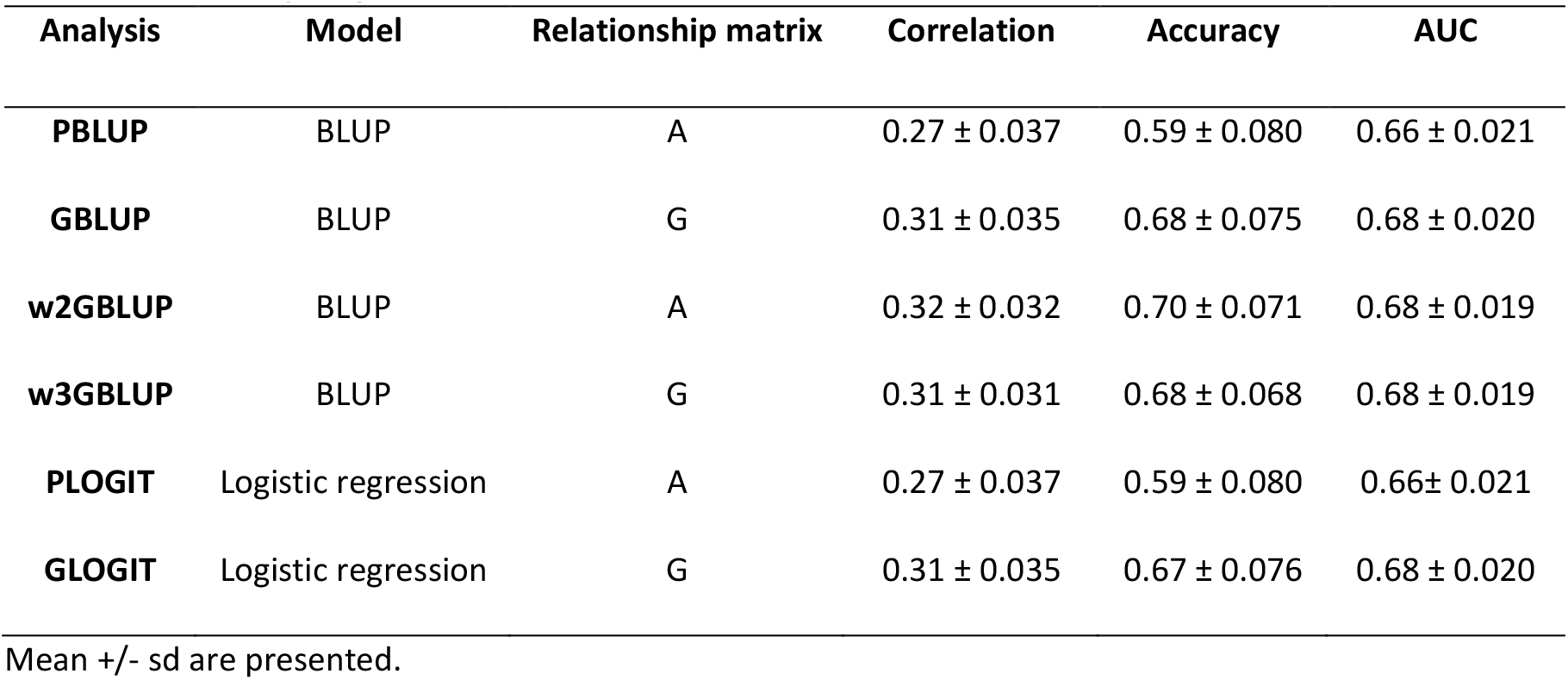
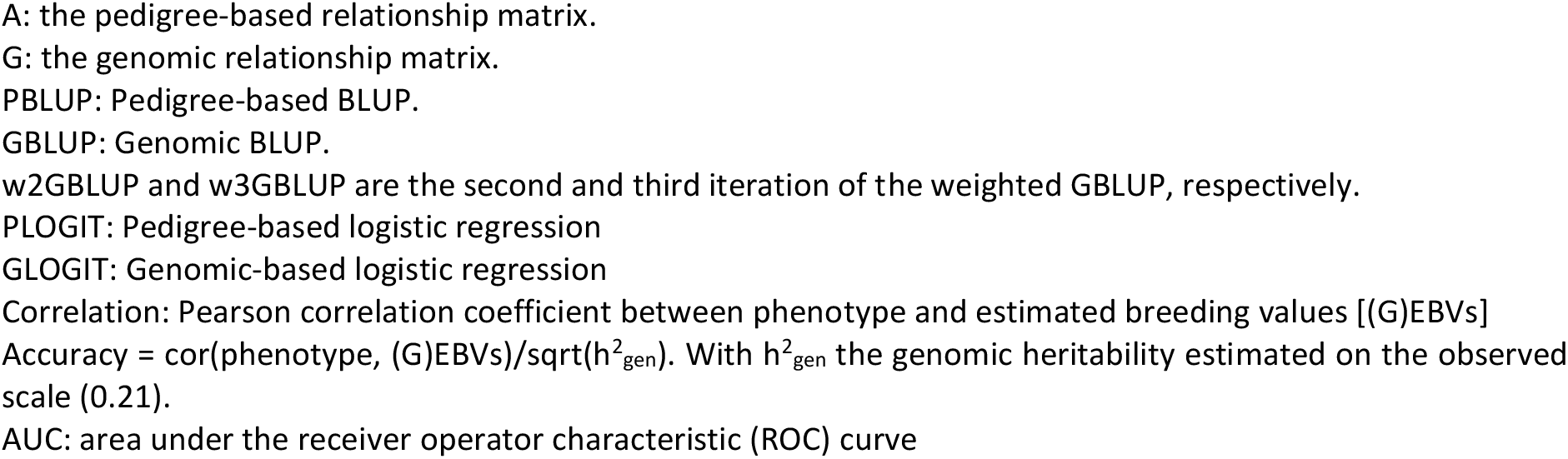
Efficiency of genomic evaluation for resistance to *F. columnare* in rainbow trout.

| Analysis | Model | Relationship matrix | Correlation | Accuracy | AUC |
| --- | --- | --- | --- | --- | --- |
| PBLUP | BLUP | A | 0.27 ± 0.037 | 0.59 ± 0.080 | 0.66 ± 0.021 |
| GBLUP | BLUP | G | 0.31 ± 0.035 | 0.68 ± 0.075 | 0.68 ± 0.020 |
| w2GBLUP | BLUP | A | 0.32 ± 0.032 | 0.70 ± 0.071 | 0.68 ± 0.019 |
| w3GBLUP | BLUP | G | 0.31 ± 0.031 | 0.68 ± 0.068 | 0.68 ± 0.019 |
| PLOGIT | Logistic regression | A | 0.27 ± 0.037 | 0.59 ± 0.080 | 0.66 ± 0.021 |
| GLOGIT | Logistic regression | G | 0.31 ± 0.035 | 0.67 ± 0.076 | 0.68 ± 0.020 |

Mean ± sd are presented.
A: the pedigree-based relationship matrix. G: the genomic relationship matrix. PBLUP: Pedigree-based BLUP. GBLUP: Genomic BLUP. w2GBLUP and w3GBLUP are the second and third iteration of the weighted GBLUP, respectively. PLOGIT: Pedigree-based logistic regression GLOGIT: Genomic-based logistic regression Correlation: Pearson correlation coefficient between phenotype and estimated breeding values [(G)EBVs] Accuracy = $\text{cor}(\text{phenotype}, (\text{G})\text{EBVs})/\sqrt{h^2_{\text{gen}}}$ . With $h^2_{\text{gen}}$ the genomic heritability estimated on the observed scale (0.21). AUC: area under the receiver operator characteristic (ROC) curve

Based on the AUC values of the ROC curves, the genomic-based relationship matrix classified the fish in the validation population with a success rate of 68% for all models, 2% better than pedigree-based relationship matrix (Table 3). No differences were observed between the linear or logistic regression models on the classifier.

## 4. Discussion

Rainbow trout, an aquaculture species of great importance worldwide, faces major threats due to infectious disease outbreaks in hatcheries and farms. Columnaris disease has become increasingly important over the past 20 years in rainbow trout production and could continue to increase as summer water temperatures rise because of global warming (Karvonen et al., 2010; Pulkkinen et al., 2010). The results presented herein show that selective breeding is a promising approach to enhance the natural resistance of broodstock, and that genomic selection may be an effective approach to increase genetic gain.

### 4.1. Resistance to *F. columnare* is moderately heritable

Resistance to *F. columnare* in this rainbow trout population was moderately heritable, with estimates ranging from of 0.18 to 0.43, pedigree-based heritability being slightly lower than genomic-based heritability. This implies that this trait can be improved by selective breeding. The heritability values estimated in the current study were in the range of previously estimated heritability (0.17-0.34) for resistance to *F. columnare* in rainbow trout populations from the USA and Norway infected in an experimental challenge (Evenhuis et al., 2015; Silva et al., 2019b). As expected, the heritability estimated on the underlying scale using a logistic regression model were significantly higher than the heritability estimated on the observed scale. When the linear estimates obtained on the observed scale were corrected as proposed by Dempster and Lerner (1950), estimates were close to the one obtained on the underlying scale with the logistic regression.

### 4.2. A major QTL on chromosome Omy3

One main genome-wide significant QTL was located on chromosome Omy3 (peak at 64.390 Mb), several less significant QTLs located on two other chromosomes (Omy12 and Omy15) as well as a polygenic background contribution. To the best of our knowledge, there is only one published study that investigated rainbow trout resistance to *F. columnare* (Silva et al., 2019a), which detected several QTLs associated with resistance in two populations, but none of the QTLs detected in the present study were reported in their study. On chromosome Omy3, the first SNP over the 5% chromosome-wide threshold was located at 38.165 Mb and the next significant SNP located at 55.715 Mb. Such a long distance between the two successive significant SNPs (17.55 Mb) may potentially reflect the presence of two QTLs (Supplementary Figure S1 and Table S1). This first SNP association may also reflect linkage disequilibrium (LD) with the peak SNP from the most significant QTL at 64.390 Mb, given that long range LD is common in rainbow trout (D’Ambrosio et al., 2019; Vallejo et al., 2018). These findings suggest that resistance to *F. columnare* in rainbow trout is oligogenic in this population, with at least one highly significant QTL and several other minor-effect QTLs and a polygenic component.

In the *Flavobacterium* genus there are two bacteria species which are responsible for diseases with similar signs and that target similar fish species. *Flavobacterium columnare*, the focus of the current study, is responsible for CD in warm waters, and *Flavobacterium psychrophilum* is responsible for cold water disease (BCWD) (Bernardet and Bowman, 2006). The symptoms of the diseases caused by these two pathogens are similar (skin lesion and septicaemia, Declercq et al., 2013; Nematollahi et al., 2003). Thus it is plausible that a certain proportion of genetic resistance mechanisms in the fish might be common between the two diseases. In the two studies on rainbow trout populations from USA and Norway, Evenhuis et al. (2015) and Silva et al. (2019b) estimated a moderate positive genetic correlation (ranging between 0.35 to 0.40) between the resistance to *F. columnare* and to *F. psychrophilum*. In addition, some of the QTLs detected in our study as associated with resistance *to F. columnare* may co-localised with previously published QTLs associated with resistance to *F. psychrophilum*. For instance, the main QTL on chromosome Omy3 (peak at 64Mb) has potentially been identified also in two previous studies on resistance to *F. psychrophilum*, after a natural outbreak in a French rainbow trout population (Fraslin et al., 2019) as well after an experimental challenge in isogenic lines of rainbow trout (Fraslin et al., 2018). Similar to our study on *F. columnare*, QTLs associated with resistance to *F. psychrophilum* in different rainbow trout populations have been previously detected on chromosomes Omy12 (Liu et al., 2015; Palti et al., 2015; Vallejo et al., 2014) and Omy15 (Fraslin et al., 2018). Even if those QTL were detected at different positions on the chromosome, they were located within wide confidence intervals and thus might be identical between the two diseases. The favourable genetic correlation for resistance to both diseases as well as the potentially common QTLs associated with resistance to *F*. *columnare* and *F. psychrophilum* on chromosomes Omy3, 12 and 15, although estimated in different rainbow trout populations with different genetic backgrounds, are encouraging for breeders. They suggest that improving the resistance to one pathogen could improve the resistance to its cold/warm counterpart.

We investigated the putative candidate genes located within a 2 Mb window around the peak SNPs using the NCBI *Oncorhynchus mykiss* Annotation Release 100 (GCF_002163495.1). Overall eleven genes involved in immune response, through the pro-inflammatory response of cytokine or the receptor-mediated endocytosis by macrophage or dendritic cells in response to bacteria activity, were located around the peak SNPs on the three chromosomes with QTLs (supplementary Table S2). Focusing on the peak SNP with the lowest *p-*value on chromosome Omy3 (located at 64,390 Mb), two genes were identified as being involved in the pro-inflammatory response of cytokine, *transforming growth factor beta receptor type-2-like (TGF-beta 2)* located between 63,826,317 and 63,853,315 bp, and an *interleukin-1 receptor type 1* (*il-1r1*) located between 65,103,069 and 65,123,204 bp. Since the QTLs detected in our study covered a large region of the chromosome, this list of putative candidate genes has to be confirmed by more studies. One approach could be to refine the QTL position using whole-genome-sequence and imputation (Fraslin et al., 2020a; Yoshida and Yáñez, 2021) to refine the list of positional candidate genes. Other approaches such as RNAseq (Marancik et al., 2015; Robledo et al., 2019, 2018; Zwollo et al., 2017) or knout-out by CRISPR/Cas9 approaches that have been recently used successfully in other fish species (Gratacap et al., 2020; Luo et al., 2022; Pavelin et al., 2021) could be used to validate their implication in the immune response to *F. columnare*.

### 4.3. Genomic evaluation increases the accuracy of breeding values

Using genomic information to predict the breeding value of fish with masked phenotype in a random cross validation scheme significantly increased the accuracy of prediction, by at least 13.6%, compared to breeding values estimated with only pedigree-based information. Both linear and logistic regression models gave similar results. The highest accuracy and highest AUC values were obtained with the weighted GBLUP approach, using the second iteration (w2GBLUP). The better performance of the weighted model compared to the standard GBLUP approach reflects the fact that, resistance to *F. columnare* in this population is controlled by a main QTL, together with a polygenic component. The interest of using a weighted approach in the presence of a main QTL have been demonstrated in various fish species for other traits of interest (Lu et al., 2020; Song and Hu, 2021; Vallejo et al., 2018). The AUC metrics obtained from the ROC curves are a way of accounting for both sensitivity (true positive rate) and specificity (true negative rate) of a test, and are a better complementary approach to accuracy of genomic prediction for binary traits (Wray et al., 2010). In the current study, the models using genomic information ((w)GBLUP and GLOGIT) were optimal for predicting the outcome of the disease. AUC values of 0.68 were obtained, in the range of what was obtained by Palaiokostas et al. (2018) for viral nervous necrosis resistance in European seabass (*Dicentrarchus labrax*), a trait with similar disease prevalence and heritability as resistance to *F. columnare* in our study (which have an impact on AUC according to Wray et al., 2010).

Prediction accuracy using genomic information in the w2GBLUP model was 18.6% higher than the pedigree-based prediction accuracy. This result along with the higher AUC values obtained when genomic information was used suggest that genomic selection is a useful approach to increase resistance to *F. columnare* in this rainbow trout population. A recently published study (Silva et al., 2019a) also concluded that genomic selection was an interesting and promising approach to increase rainbow trout resistance to *F. columnare* due to the polygenic architecture of the trait. They observed an improved prediction accuracy of about 40% when using genomic models compared to pedigree-based models. Those results along with the one of the current study are promising since they confirm the major benefit of using genomic selection to improve resistance to *F. columnare* in different rainbow trout populations, with different genetic architecture of resistance. It should be noted that diseases like *F. columnare* infect small fish well before they can be individually tagged, and hence genotyping is needed to establish relationships between individuals. Therefore, the cost-efficiency of genomic selection may be reasonable, since genotyping is routinely performed anyway, even if at lower density than the SNP array.

### 4.4. Phenotyping for resistance after a natural disease outbreak

In our study, we took advantage of a natural outbreak of columnaris disease to detect QTL associated with resistance to the pathogen agent responsible of the disease and to estimate the potential of selective breeding to increase the resistance of this rainbow trout population. Experimental challenges are usually used for disease resistance studies since they allow a more controlled experiment, knowing the time when the fish was infected and when it died or sometimes showed signs of infection (Fraslin et al., 2020b; Ødegård et al., 2011; Robinson et al., 2017; Saura et al., 2019). However, experimental challenge requires advanced knowledge of the bacteria in order to isolate and replicate it while still keeping the infectivity high enough to induce mortality in an infectious challenge. Experimental pathogen challenge usually needs to be performed in different facilities, under strict controls, and thus could be expensive to set up within a breeding programme. Furthermore the results obtained in a controlled environment would potentially need to be validated in a field setting before being implemented in a commercial breeding programme (Wiens et al., 2018). Field or natural outbreak data can provide very valuable phenotypes for disease resistance, as the fish are exposed to realistic commercial conditions. In various fish species resistance measured after an natural disease outbreak has been successfully used to estimate genetic parameters (Bangera et al., 2014; Barría et al., 2020; Lillehammer et al., 2013) and detect QTL associated with resistance (Barría et al., 2021; Fraslin et al., 2018; Houston et al., 2008). However, they also have disadvantages such as being unpredictable and the uncertainty of which specific pathogens are responsible for the observed disease or mortality. Furthermore, in the current study, the fish had to be treated for the disease. Thus, the fish that were considered as resistant fish for being still alive at the end of the 3 months rearing periods might not all be truly resistant fish but fish that were very slow to develop disease symptoms, or by chance were not in contact with the bacteria before the treatment, or fish that were infected but cured by the treatment they received. Thus, the resistance phenotype measured in our study may not be the true resistance to the pathogen as usually defined in disease resistance studies (Fraslin et al., 2020b; Robinson et al., 2017). However, the co-location between the QTL detected in our study and previously published QTLs associated with resistance to *F. psychrophilum* (mainly on Omy3) suggests that resistance measured in the current study is indeed an appropriate measure of genetic resistance. Furthermore, the results of the current study suggest that phenotyping of resistance after a natural outbreak can be performed as part of the normal rearing process with little effort (collecting the dead fish prior to the treatment) which is potentially less time consuming and less expensive than designing an experimental challenge on siblings of the breeding candidates.

## 5. Conclusion

In the current study, a moderate heritability of resistance to *Flavobacterium columnare* was estimated in a rainbow trout population after natural disease outbreak. Resistance was controlled by a major QTL, located on chromosome Omy3, together with various smaller QTLs and a polygenic background contribution. Finally, genomic selection was shown to potentially be an efficient solution to improve genetic resistance by selective breeding, giving approximately 14% higher accuracy of breeding values than pedigree approaches.

## Acknowledgment

The skilled staff of Savon Taimen Oy and Hanka Taimen Oy are thanked for their expertise in data collection and fish rearing.

## Author contribution

**AK, RDH AN** and **HK** were responsible for the concept and design of this work. **AK** and **RDH** were responsible for the supervision of the work and funding acquisition. **AK, AN** and **HK** were responsible of fish management and data collection. **CF** performed bioinformatics and statistical analyses. **CF, RDH** and **AK** drafted the manuscript. All authors read and approved the final manuscript.

## Funding

This work is part of the AquaIMPACT project and was supported by the European Union’s Horizon 2020 research and innovation programme under the grant agreement No 818367. The Roslin Institute received BBSRC Institute Strategic Program funding (BB/P013732/1, BB/P013740/1, BB/P013759/1).

## Supplementary Tables

**Table S1.** All SNPs that are significant (5% chromosome wide level, Bonferroni correction), in red the peak SNPs.

| Omy | SNP name | Position (bp) | a (SNP additive effect) | se | p-value | -log10(p-value) |
| --- | --- | --- | --- | --- | --- | --- |
| 3 | AX-89921627 | 38,165,148 | -7.08E-02 | 1.65E-02 | 1.84E-05 | 4.7341 |
| 3 | AX-89963942 | 55,714,839 | 8.40E-02 | 2.03E-02 | 3.67E-05 | 4.4357 |
| 3 | AX-89960286 | 63,823,874 | -7.83E-02 | 1.72E-02 | 5.37E-06 | 5.2700 |
| 3 | AX-89974385 | 64,017,036 | 9.37E-02 | 1.63E-02 | 1.01E-08 | 7.9967 |
| 3 | AX-89959477 | 64,035,510 | -7.19E-02 | 1.64E-02 | 1.10E-05 | 4.9580 |
| 3 | AX-89949124 | 64,057,117 | -7.91E-02 | 1.90E-02 | 3.13E-05 | 4.5038 |
| 3 | AX-89952021 | 64,375,741 | 9.85E-02 | 1.76E-02 | 2.36E-08 | 7.6269 |
| 3 | AX-89933339 | 64,390,419 | -1.10E-01 | 1.83E-02 | 1.69E-09 | 8.7731 |
| 3 | AX-89959970 | 64,778,731 | -1.02E-01 | 1.92E-02 | 1.17E-07 | 6.9301 |
| 3 | AX-89973040 | 64,779,737 | 9.36E-02 | 2.18E-02 | 1.76E-05 | 4.7535 |
| 3 | AX-89919724 | 64,781,151 | -8.68E-02 | 1.86E-02 | 2.93E-06 | 5.5334 |
| 3 | AX-89968862 | 64,785,727 | 9.24E-02 | 2.27E-02 | 4.54E-05 | 4.3429 |
| 3 | AX-89944666 | 65,224,088 | -1.11E-01 | 2.17E-02 | 3.30E-07 | 6.4814 |
| 3 | AX-89973043 | 65,401,327 | -1.07E-01 | 2.21E-02 | 1.37E-06 | 5.8639 |
| 3 | AX-89952255 | 65,429,923 | -9.98E-02 | 2.09E-02 | 1.79E-06 | 5.7480 |
| 3 | AX-89976032 | 65,733,733 | 8.08E-02 | 1.75E-02 | 4.06E-06 | 5.3914 |
| 3 | AX-89967905 | 68,232,938 | 8.20E-02 | 2.01E-02 | 4.54E-05 | 4.3427 |
| 3 | AX-89939438 | 70,211,721 | -1.02E-01 | 2.45E-02 | 3.17E-05 | 4.4993 |
| 3 | AX-89946264 | 70,261,448 | -1.09E-01 | 2.32E-02 | 2.44E-06 | 5.6128 |
| 3 | AX-89929491 | 70,982,024 | 1.10E-01 | 2.18E-02 | 4.34E-07 | 6.3628 |
| 3 | AX-89924934 | 75,773,376 | -1.19E-01 | 2.53E-02 | 2.83E-06 | 5.5489 |
| 3 | AX-89945270 | 76,567,992 | -1.21E-01 | 2.59E-02 | 2.89E-06 | 5.5395 |
| 3 | AX-89933590 | 79,566,898 | -1.44E-01 | 3.54E-02 | 4.73E-05 | 4.3249 |
| 12 | AX-89951575 | 5,316,265 | -7.28E-02 | 1.72E-02 | 2.30E-05 | 4.6377 |
| 15 | AX-89927482 | 13,366,111 | 6.69E-02 | 1.52E-02 | 1.06E-05 | 4.9756 |
| 15 | AX-89956130 | 41,172,606 | 9.30E-02 | 2.13E-02 | 1.25E-05 | 4.9015 |
| 15 | AX-89967415 | 46,205,286 | -7.01E-02 | 1.69E-02 | 3.42E-05 | 4.4664 |
| 15 | AX-89942287 | 47,018,101 | -7.03E-02 | 1.71E-02 | 4.06E-05 | 4.3911 |

**Table S2.**
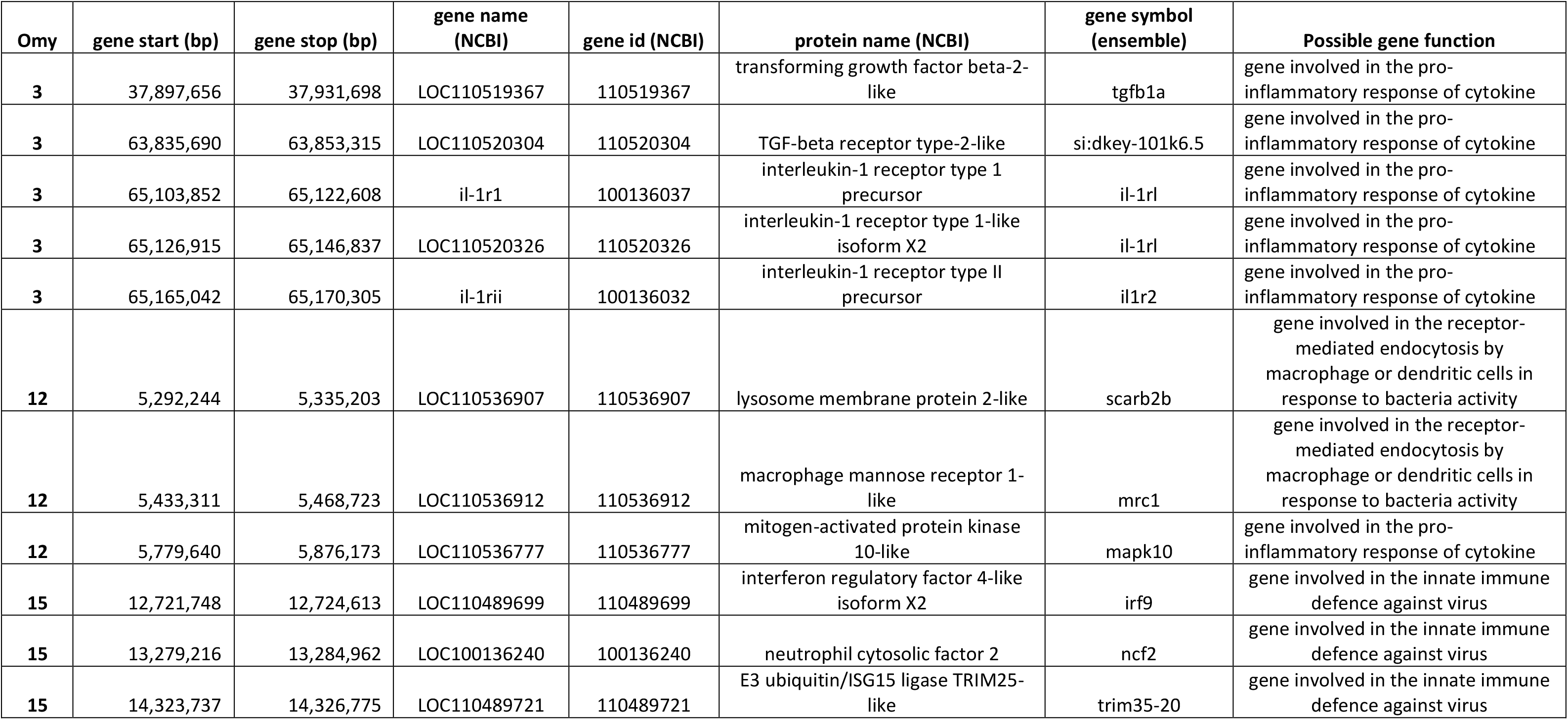
All genes located in 2 Mb around the peak SNPs.

---

**Supplementary Figure S1.**
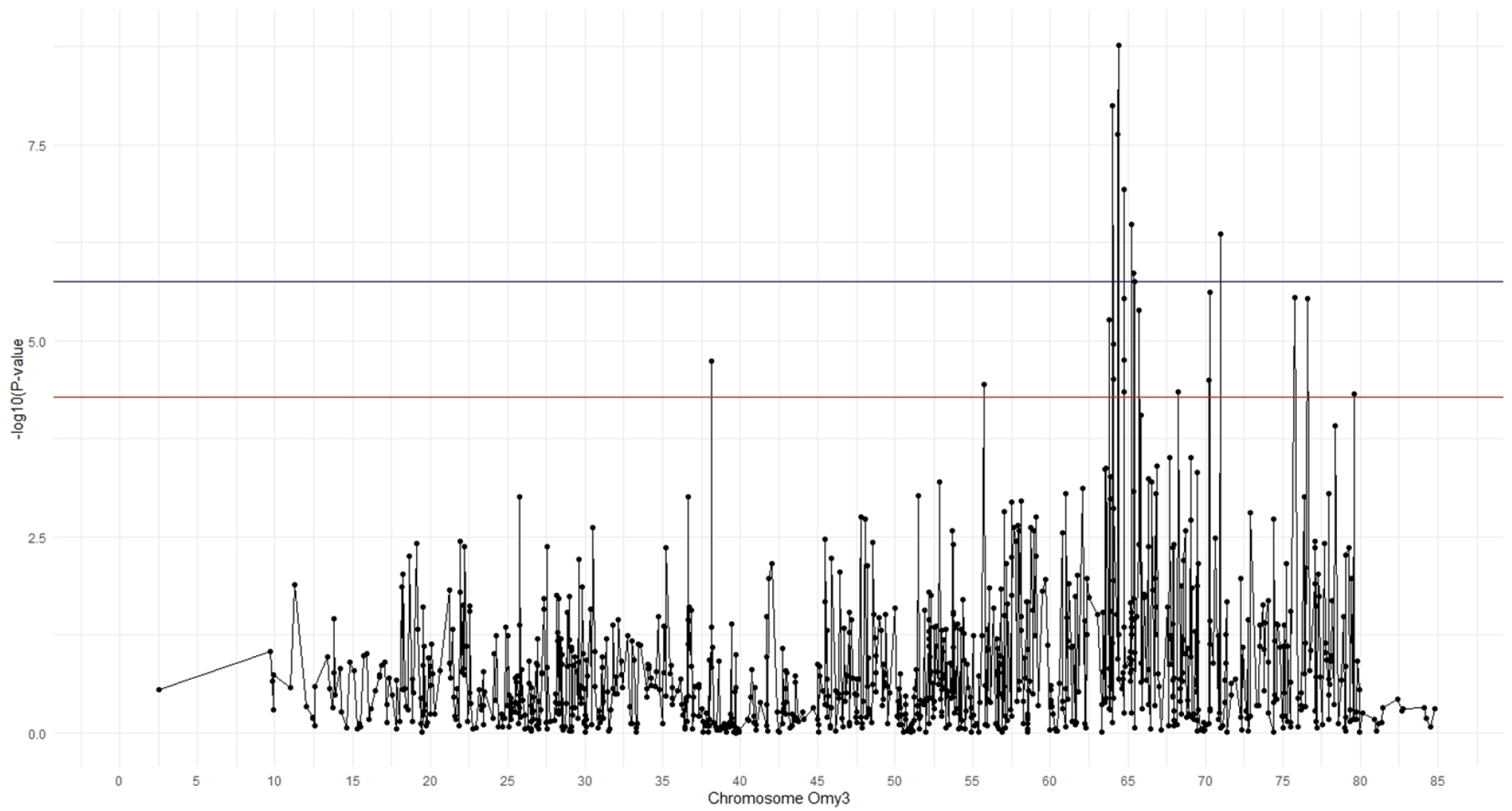
Manhattan plot of the chromosome Omy3. The dark blue line is the 5% significance threshold at the genome-wide level, the red line is the 5% suggestive threshold at the chromosome-wide level.

## Notes

### Competing Interest Statement

The authors have declared no competing interest.

